# Inherited chromosomally integrated human herpesvirus 6 genomes are ancient, intact and potentially able to reactivate from telomeres

**DOI:** 10.1101/166041

**Authors:** Enjie Zhang, Adam J Bell, Gavin S Wilkie, Nicolás M Suárez, Chiara Batini, Colin D Veal, Isaac Armendáriz-Castillo, Rita Neumann, Victoria E Cotton, Yan Huang, David J Porteous, Ruth F Jarrett, Andrew J Davison, Nicola J Royle

**Affiliations:** Department of Genetics, University of Leicester, Leicester, UK; MRC-University of Glasgow Centre for Virus Research, Glasgow, UK; Department of Health Sciences, University of Leicester, Leicester, UK; Generation Scotland, Centre for Genomic and Experimental Medicine, Institute of Genetics and Molecular Medicine, University of Edinburgh, UK

**Keywords:** Human herpesvirus 6, telomere, integration, ciHHV-6, molecular dating, Generation Scotland

## Abstract

Human herpesviruses 6A and 6B (HHV6-A and HHV-6B; species *Human herpesvirus 6A* and *Human herpesvirus 6B*) have the capacity to integrate into telomeres, the essential capping structures of chromosomes that play roles in cancer and ageing. About 1% of people worldwide are carriers of chromosomally integrated HHV-6 (ciHHV-6), which is inherited as a genetic trait. Understanding the consequences of integration for the evolution of the viral genome, for the telomere and for the risk of disease associated with carrier status is hampered by a lack of knowledge about ciHHV-6 genomes. Here, we report an analysis of 28 ciHHV-6 genomes and show that they are significantly divergent from the few modern non-integrated HHV-6 strains for which complete sequences are currently available. In addition ciHHV-6B genomes in Europeans are more closely related to each other than to ciHHV-6B genomes from China and Pakistan, suggesting regional variation of the trait. Remarkably, at least one group of European ciHHV-6B carriers has inherited the same ciHHV-6B genome, integrated in the same telomere allele, from a common ancestor estimated to have existed 24,500 ±10,600 years ago. Despite the antiquity of some, and possibly most, germline HHV-6 integrations, the majority of ciHHV-6B (95%) and ciHHV-6A (72%) genomes contain a full set of intact viral genes and therefore appear to have the capacity for viral gene expression and full reactivation.

**IMPORTANCE:** Inheritance of HHV-6A or HHV-6B integrated into a telomere occurs at a low frequency in most populations studied to date but its characteristics are poorly understood. However, stratification of ciHHV-6 carriers in modern populations due to common ancestry is an important consideration for genome-wide association studies that aim to identify disease risks for these people. Here we present full sequence analysis of 28 ciHHV-6 genomes and show that ciHHV-6B in many carriers with European ancestry most likely originated from ancient integration events in a small number of ancestors. We propose that ancient ancestral origins for ciHHV-6A and ciHHV-6B are also likely in other populations. Moreover, despite their antiquity, all of the ciHHV-6 genomes appear to retain the capacity to express viral genes and most are predicted to be capable of full viral reactivation. These discoveries represent potentially important considerations in immune-compromised patients, in particular in organ transplantation and in stem cell therapy.

## INTRODUCTION

Given the complex roles that human telomeres play in cancer initiation and progression and in ageing (1, 2), it is remarkable that the genomes of human herpesviruses 6A and 6B (HHV6-A and HHV-6B; species *Human herpesvirus 6A* and *Human herpesvirus 6B*) can integrate and persist within them (3). Human telomeres comprise double-stranded DNA primarily composed of variable lengths of (TTAGGG)_n_ repeats and terminated by a 50-300 nucleotide (nt) 3’ single-strand extension of the G-rich strand. Telomeres, bound to a six-protein complex called shelterin, cap the ends of chromosomes and prevent inappropriate double-strand break repair. They also provide a solution to the ‘end replication problem’ via the enzyme telomerase (4-6).

The double-stranded DNA genomes of HHV-6A and HHV-6B consist of a long unique region (U; 143-145 kb) encoding many functional open reading frames (ORFs U2-U100), flanked by identical left and right direct repeats (DR_L_ and DR_R_; 8-10 kb) encoding two ORFs (DR1 and DR6). Each DR also contains near its ends two variable regions of telomere-like repeat arrays (T1 and T2) (7, 8), terminated by the viral genome packaging sequences (PAC1 and PAC2, respectively) (9, 10). Telomeric integration by HHV-6A or HHV-6B (yielding chromosomally integrated HHV-6, ciHHV-6) results in loss of the terminal PAC2 sequence at the fusion point between the telomere and DR_R_-T2 (11) and loss of the DR_L_-PAC1 sequence at the other end of the integrated viral genome when the DR_L_-T1 degenerate telomere-like repeat region becomes part of a newly formed telomere (Figure 1A, (12)).

**Figure 1.**
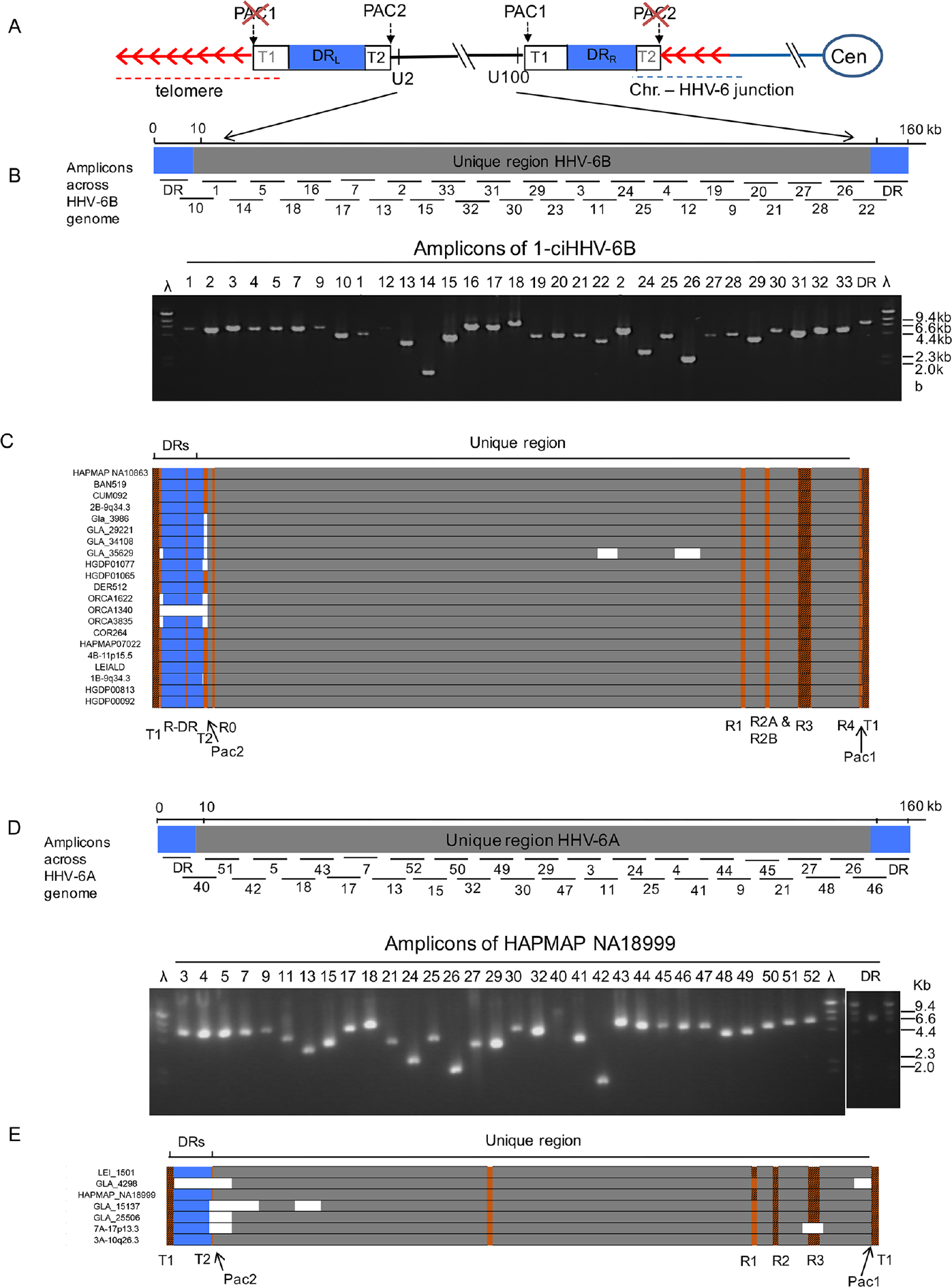
Approach to sequencing ciHHV-6 genomes. (A) Diagram showing the organisation of the HHV-6 genome following integration of a single full-length copy into a telomere. Chromosome and centromere (Cen) are shown by blue lines and an oval. The telomere repeats are shown by red arrows. The telomere, encompassing DR_L_-T1, is shown by a red dashed line. The junction between the chromosome and HHV-6 genome, encompassing telomere repeats and DR_R_-T2, is shown by a dashed blue line. DR_L_ and DR_R_ are shown as blue boxes. (B) Distribution of numbered PCR amplicons across the HHV-6B genome and an example gel of PCR products generated from 1-ciHHV-6B. (C) Sequence coverage for individual ciHHV-6B genomes. Each ciHHV-6B genome is shown with a single DR (blue box) that was covered by amplicons from DR_L_ and DR_R_ and with U (grey box). Gaps in the coverage caused by loss of individual amplicons at the amplicon-pooling stage are shown in white. Tandem repeat regions that were fully sequenced by either Illumina NGS or by the Sanger method are shown in orange. Tandem repeat regions (e.g. T1 and R3 in HHV-6B) that were too long to be sequenced fully are shown as hashed-brown boxes. (D) Distribution of numbered PCR amplicons across the HHV-6A genome and an example gel of products generated from HAPMAP NA18999. (E). Sequence coverage for each ciHHV-6A genomes, using the same colour coding as in (C).

Once the HHV-6 genome has integrated in the germline it can be passed from parent to child, behaving essentially as a Mendelian trait (inherited ciHHV-6) (13-16). The telomere carrying the ciHHV-6 genome shows instability in somatic cells, which can result in the partial or complete release of the viral genome as circular DNA (12, 17, 18). This could represent the first step towards viral reactivation, and in this respect telomeric integration may be a form of HHV-6 latency. To date, reactivation of ciHHV-6 has been demonstrated in two settings: first, in a child with X-linked severe combined immunodeficiency who was also a carrier of inherited ciHHV-6A (19); and second, upon transplacental transmission from two ciHHV-6 carrier mothers to their non-carrier babies (20). Recently, it has been shown that ciHHV-6 carriers bear an increased risk of angina pectoris (21), although it is not known whether this arises from viral reactivation, a deleterious effect on the telomere carrying the viral genome, or some other mechanism.

A small proportion of people worldwide are carriers of inherited ciHHV-6A or -6B but very little is known about the HHV-6 genomes that they harbor, although that may influence any associated disease risk. To investigate ciHHV-6 genomic diversity and evolution, the frequency of independent germline integrations, and the potential functionality of the integrated viral genomes, we analysed 28 ciHHV-6 genomes. We discovered that ciHHV-6 genomes are more similar to one another than to the few sequenced reference HHV-6 genomes from non-integrated viruses. This is particularly marked among the ciHHV-6B genomes from Europeans. We also found that a subset of ciHHV-6B carriers from England, Orkney and Sardinia are most likely descendents from a single ancient ancestor. Despite the apparent antiquity of some, possibly most, ciHHV-6 genomes, we concluded that the majority contain a full set of intact HHV-6 genes and therefore in principle retain the capacity to generate viable viruses.

## MATERIAL AND METHODS

### Population screening to identify ciHHV-6 carriers

ciHHV-6 carriers were identified by screening a variety of DNA sample collections of individuals from across the world, using PCR assays to detect either U11, U18, DR5 (HHV-6A) or DR7 (HHV-6B) (12), or U7, DR1, DR6A or DR6B ((22) and manuscript in preparation). DR5, DR6A, DR6B and DR7 correspond to ORFs in the original annotation of the HHV-6A genome (GenBank accession X83413 (23)), but DR5 is in a non-coding region of the genome, and DR6A, DR6B and DR7 are in exons of DR6 in the reannotation used (RefSeq accession NC_001664). From the populations screened, 58 samples with ciHHV-6 among 3875 individuals were identified (Supplementary Table 1). The number of individuals screened in most populations was small and therefore cannot be used to give an accurate estimate of ciHHV-6A or -B frequencies, although a larger number of ciHHV-6B-positive samples was identified overall. The frequency of ciHHV-6B carriers in Orkney (1.9%), a collection of islands off the north coast of Scotland, is higher than that reported from England (24). Screening of the Generation Scotland: Scottish Family Health Study (GS:SFHS) will be described elsewhere (RFJ manuscript in preparation). Ethical approval for the GS:SFHS cohort was obtained from the Tayside Committee on Medical Research Ethics (on behalf of the National Health Service).

### Generation of overlapping amplicons and sequencing

The 32 primer pairs used to generate overlapping amplicons from ciHHV-6A genomes, and the PCR conditions employed, were reported previously (18). The primer pairs used to amplify ciHHV-6B genomes were based on conserved sequences from the HHV-6B non-integrated HST and Z29 strains (Genbank accessions AB021506.1 and AF157706 respectively; (9, 25). The primer sequences are shown in Supplementary Table S2. The amplicons from each sample were pooled in equimolar proportions and then sequenced using the Illumina MiSeq or IonTorrent (Life Technologies) next-generation sequencing platforms, as described previously (18). Some sequences were verified by using Sanger dideoxy chain termination sequencing on PCR-amplified products.

### Assembly and analysis of DNA sequence data

DNA sequence data were processed essentially as described previously (18), except that SPAdes v. 3.5.0 (26) was used for *de novo* assembly into contigs, ABACAS v. 1.3.1 (27) was used to order contigs, and Gapfiller v. 1-11 (28) was used to fill gaps between contigs. The integrity of the sequences was verified by aligning them against the read data using BWA v. 0.6.2-r126 and visualizing the alignments as BAM files using Tablet v. 1.13.08.05. Nucleotide substitutions, indels and repeat regions were also verified by manual analysis using IGV v. 2.3 (http://software.broadinstitute.org/software/igv/home).

Alignments of the seven iciHHV-6A genomes with the three published HHV-6A genomes from non-integrated strains U1102, GS and AJ (23, 29-31) and alignment of the 21 iciHHV-6B genomes with the two previously published HHV-6B genomes from non-integrated viruses HST and Z29 (9, 25) were carried out by using Gap4 (32). Variation across the iciHHV-6 genomes was studied by a combination of manual inspection and automated analysis by using an in-house Perl script. The script performed a sliding window count of substitutions using the aligned GAP4 files, reporting the count according to the mid-point of the window. For analysis across the genome, the window size was 1 kb and the step size was 1 nucleotide. For analysis of individual ORFs, a file with a list of annotated positions was generated.

Phylogenetic analyses were carried out using two different methods. Maximum likelihood trees were built by using the Maximum Composite Likelihood model (MEGA6.0) and bootstrap values obtained with 2000 replications. Model selection was carried out for HHV-6A and HHV-6B separately, and the substitution model with the lowest Bayesian information criterion was selected (the Tamura 3-parameter model (33) for HHV-6B and the Hasegawa-Kishino-Yano model for HHV-6A). Median-joining networks were built by using Network 5.0 (www.fluxus-engineering.com) with default parameters. Sites with missing data were excluded from all phylogenetic analyses for both HHV-6A and HHV-6B. The number of positions analysed for HHV-6B was 130412, and that for HHV-6A was 117900. Time to the most recent common ancestor (TMRCA) was calculated by using rho as implemented in Network 5.0. Rho values were transformed into time values by using the accepted mutation rate for the human genome, 0.5E-09 substitutions per bp per year (34), scaled to the number of sites analysed.

### Comparison of tandem repeat regions

The copy number of repeat units in the DR-R, R0, R1, R2, R3 and R4 tandem repeat regions (9, 25) were determined by manual inspection of the individual BAM files generated for each sequenced ciHHV-6 genome, with verification by checking the sequence alignments generated using Gap4. The numbers of copies of TTAGGG in each DR_L_-T2 region was determined from PCR amplicons generated using the DR8F and UDL6R primers (Supplementary Table S2). Each amplicon was purified using a Zymoclean™ gel DNA recovery kit, and then sequenced using the Sanger dideoxy chain termination method. The sequence data were analysed using MacVector software. Variation at the (CA)_n_ repeat array located immediately adjacent to T1 in HHV-6B was investigated in DR_L_ specifically by reamplification of single telomere length analysis (STELA) products, using the primers DR1R and TJ1F. The short amplicons were purified and sequenced as above and compared with the same sequence in the reference HST and Z29 genomes.

### Analysis of DR_R_-T1 region by TVR-PCR

The DR_R_-T1 regions from ciHHV-6B positive samples were amplified using the primers U100Fw2 and DR1R. Telomere variant repeat mapping by PCR (TVR-PCR) was conducted on each of these amplicons essentially as described before (35, 36) but using an end-labeled primer, HHV-6B-UDR5F and the unlabeled TAG-TELWRev. The TELWRev primer anneals to TTAGGG repeats allowing amplification of products that differ in length depending on the location of the TTAGGG repeat with respect to the flanking primer (HHV-6B-UDR5F). The labeled amplicons from the T1 region were size separated in a 6% denaturing polyacrylamide gel.

### Analysis of HHV-6 ORFs

The frequency of nucleotide substitutions in each ORF was determined by a combination of manual inspection and automated analysis using a Perl script, as described above. The DNA sequences of each of the 86 HHV-6B ORFs from the 21 ciHHV-6B genomes were aligned to identify and compare the number of synonymous and non-synonymous codon changes within and among genes. In addition the predicted amino acid sequences for each gene in the 21 ciHHV-6B genomes were aligned to confirm the number of non-synonymous changes.

### Characterisation of chromosome-ciHHV-6 junctions

The junctions between the chromosome and the ciHHV-6 genome were isolated by PCR amplification using various primers that anneal to subterminal regions of a variety of human chromosomes in combination with the DR8F primer. The amplicons were purified as described above and sequenced by using the Sanger method with a variety of primers (Supplementary Table S2). The number of repeats present in each junction fragment and the interspersion of TTAGGG repeats with degenerate repeats was determined by manual inspection using the MacVector software.

## RESULTS

### Selection of ciHHV-6 carriers and sequence analysis of viral genomes

To investigate sequence variation among ciHHV-6 genomes, 28 samples were selected for analysis: seven with ciHHV-6A (including LEI-1501 (18)) and 21 with ciHHV-6B (Table 1). The selected samples were identified in the various populations screened (Supplementary Table S1), and included additional individuals from the London area (16), Scotland and the north of England (22), the Leicester area of England (18) and the GS: SFHS (RJF manuscript in preparation). The chromosomal location of ciHHV-6 genomes, determined by fluorescent *in situ* hybridisation (FISH), was available for some samples (16, 18). For other samples the junction between the viral DR8 sequence (a non-coding region near one end of DR) and the chromosome subtelomeric region was isolated by PCR and sequenced (discussed below). Integration of each ciHHV-6 genome was confirmed by detection of a telomere at DR_L_-T1 using STELA (12), or by detection of at least one copy per cell using droplet digital PCR (22).

**Table 1.** Samples from individuals with ciHHV-6 selected for viral genome sequencing.

| Sample | ciHHV-6 species | Integration Site <sup>a</sup> | Population or Country |
| --- | --- | --- | --- |
| LEI-1501 <sup>b</sup> | A | 19q | Leicester area (England) |
| HAPMAP NA18999 | A |  | Japan |
| 3A-10q26.3 <sup>c</sup> | A | 10q26.3 & junction isolated by PCR | South-East England |
| GLA_4298 <sup>d</sup> | A |  | Newcastle (England) |
| GLA_15137 <sup>e</sup> | A |  | Scotland |
| GLA_25506 <sup>e</sup> | A |  | Scotland |
| 7A-17p13.3 <sup>c</sup> | A | 17p13.3 | South-East England |
| HGDP00092 | B |  | Balochi (Pakistan) |
| HGDP00813 | B |  | Han (China) |
| HGDP01065 | B | Junction isolated by PCR | Sardinia (Italy) |
| HGDP01077 | B | Junction isolated by PCR | Sardinia (Italy) |
| HAPMAP NA07022 (CEPH 1340.11) | B |  | Utah Mormon (North European) |
| HAPMAP NA10863 (CEPH-1375.02) | B |  | Utah Mormon (North European) |
| 4B-11p15.5 <sup>c</sup> | B | 11p15.5 | Southern East England |
| BAN519 <sup>f</sup> | B |  | Banff (Scotland) |
| COR264 <sup>f</sup> | B |  | Cornwall (England) |
| CUM082 <sup>f</sup> | B |  | Cumbria (England) |
| DER512 <sup>f</sup> | B | Junction isolated by PCR | Derbyshire (England) |
| 2B-9q34.3 <sup>c</sup> | B | 9q34.3 | South-East England |
| 1-ciHHV-6B <sup>c</sup> | B | Junction isolated by PCR | South-East England |
| LEI-ALD | B |  | Leicester area (England) |
| ORCA1622 | B |  | Orkney |
| ORCA1340 | B |  | Orkney |
| ORCA3835 | B |  | Orkney |
| GLA_3986 <sup>d</sup> | B |  | Newcastle (England) |
| GLA_29221 <sup>e</sup> | B |  | Scotland |
| GLA_34108 <sup>e</sup> | B |  | Scotland |
| GLA_35629 <sup>e</sup> | B |  | Scotland |

<sup>a</sup> Determined by FISH or amplification of chromosome-ciHHV-6 junctions by PCR; <sup>b</sup> LEI-1501 described in (18); <sup>c</sup> ciHHV-6 carriers describe in (16); <sup>d</sup> ciHHV-6 carriers previously described in (22); <sup>e</sup> ciHHV-6 carriers identified in the GS: SFHS; <sup>f</sup> Samples from the Population of British Isles study (42).

Each viral genome from ciHHV-6 carriers was sequenced from pooled PCR amplicons (12, 18). Full sets of HHV-6 amplicons were readily generated (Fig. 1 and Supplementary Table S2), demonstrating the robustness of this approach for enriching HHV-6 sequences from ciHHV-6 carriers. The HHV-6 amplicons generated from each carrier had the expected sizes, with variation only in amplicons encompassing repetitive regions (e.g. the DR_R_-T1 region of degenerate telomere-like repeats). This observation indicated that all of the ciHHV-6 genomes are essentially intact, with the exception of the terminal DR_R_-PAC2 and DR_L_-PAC1 sequences lost during integration (Fig. 1A) (11, 12).

The ciHHV-6 genome sequences were determined by short-read next-generation sequencing (NGS) with some verification by the Sanger method. De novo assemblies of each genome were generated with few gaps (Fig. 1). The finished sequences were annotated and deposited in GenBank under accession numbers KY316030-KY316056. The ciHHV-6A genome reported previously by us was included in these analyses (LEI-1501, KT355575; (18)).

### Sequence similarity is greater among ciHHV-6 genomes than to non-integrated HHV-6 genomes

Nucleotide substitution frequencies were analysed across the DR and U regions of the HHV-6B genome (excluding the tandem repeat regions R-DR, R0, R1, R2, R3 and R4, see Fig. 1, (9, 25)) for each sequenced ciHHV-6B genome in comparison with the two available HHV-6B reference genomes from non-integrated strains (HST from Japan, GenBank accession AB021506, (25) and Z29 from Democratic Republic of Congo (D.R.Congo), GenBank accession AF157706, (9)). The ciHHV-6B genomes show different patterns of variation from the reference genomes, with greater divergence from strain Z29 in the distal portion of the U region (120-150 kb) and across DR (1-8 kb), reaching a maximum of 35 substitutions per kb in these regions (Fig. 2A). Overall, there is less divergence from strain HST, although the frequency of substitutions is higher in part of the U region (45-64 kb) compared to strain Z29. To assess sequence variation among the ciHHV-6B genomes, comparisons were made using the genome in HAPMAP NA10863 (CEPH1375.02) as a reference. The substitution frequency is considerably less across the viral genomes for 18/20 of the ciHHV-6B genomes from individuals with European ancestry, indicating greater similarity among them. Notably, the other two ciHHV-6B genomes that showed a higher substitution frequency in this comparison were in individuals from Pakistan and China, HGDP00092 and HGDP00813, respectively (Fig. 2A).

**Figure 2.**
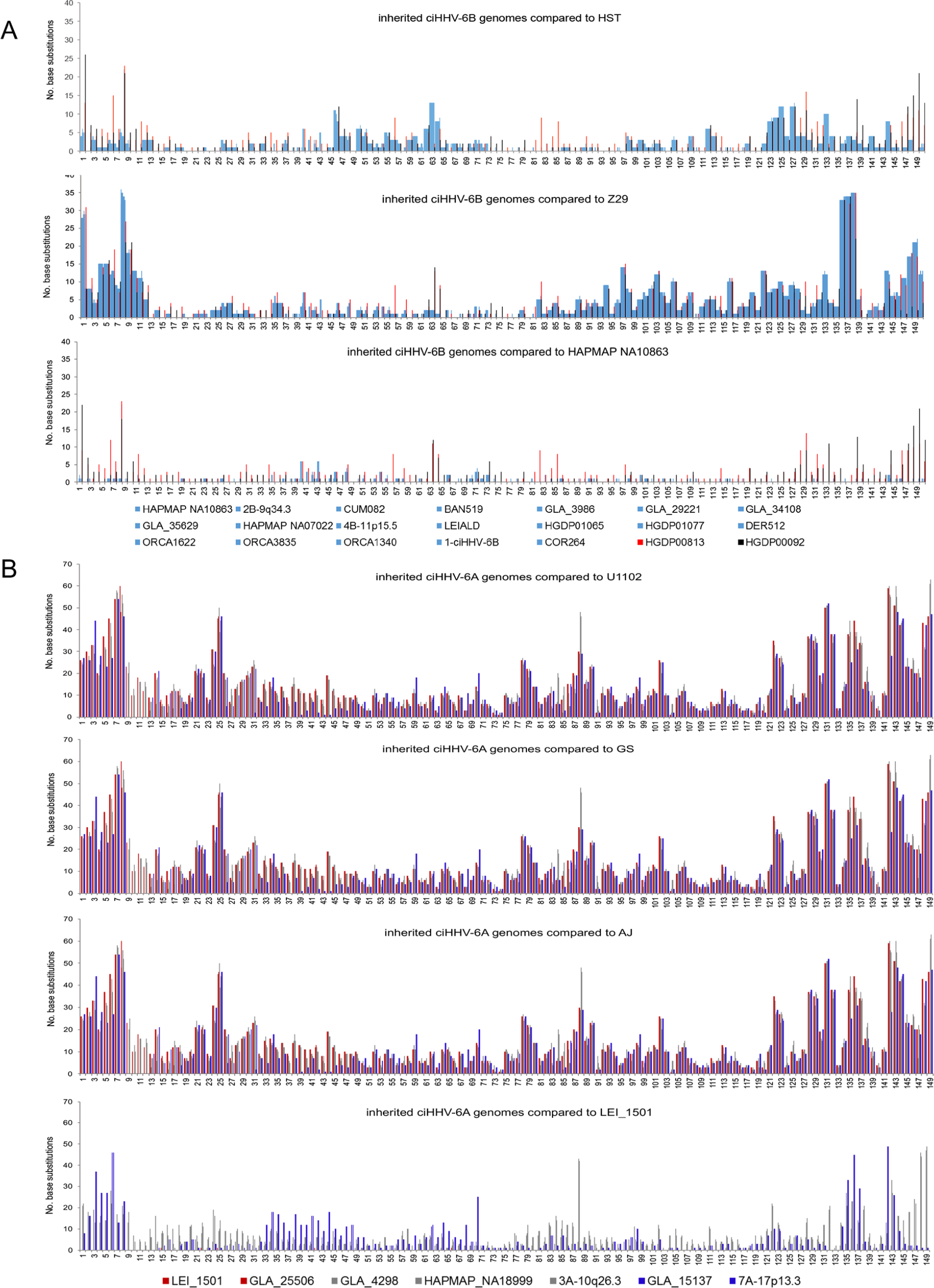
Frequency of nucleotide substitutions in ciHHV-6 genomes compared to reference viral genomes. (A) Graphs showing the number of substitutions in 1 kb windows for each of the 21 ciHHV-6B genomes in comparison with the HHV-6B strain HST (Japan) and Z29 (D.R.Congo) genomes (top and middle panels, respectively) and the ciHHV-6B genome from HAPMAP NA10863 (bottom panel). The colour-coded key shows that ciHHV-6B genomes from individuals with European ancestry are represented as light blue lines; ciHHV-6B in HGDP00813 (China), red lines; ciHHV-6B in HGDP00092 (Pakistan), black lines. (B) Graphs showing the number of substitutions in 1 kb windows for each of the 7 ciHHV-6A genomes in comparison with the HHV-6A strain U1102 (Uganda), GS (USA) and AJ (Gambia) genomes (top and two middle panels) and the ciHHV-6A genome in LEI-1501 (bottom panel). The colour-coded key distinguishes each of the ciHHV-6A genomes. The x-axes in all the graphs show the HHV-6B and -6A genomes with a single DR (0-8kb) followed by U (9-150kb) as shown in Figure 1C and 1E. Variation within the tandem repeat regions is not shown in these graphs.

Nucleotide substitution frequencies were also analysed across each of the seven ciHHV-6A genomes in comparison with three non-integrated HHV-6A reference genomes (strain U1102 from Uganda (37) (accession X83413, (23)); strain GS from the USA (accessions KC465951.1 (GS1) and KJ123690.1 (GS2) (29, 30)) and strain AJ from the Gambia (accession KP257584.1 (31)). This analysis shows that the ciHHV-6A genomes have similar levels of divergence from each reference genome from non-integrated HHV-6A and that divergence is highest across DR and the distal part of U (120-149 kb) (Fig. 2B). Comparisons with the ciHHV-6A LEI-1501 genome (18) as a reference, also showed greater similarity among the ciHHV-6A genomes although the substitution frequencies are higher than among European ciHHV-6B genomes, indicating greater diversity among the ciHHV-6A genomes sequenced here (38). Notably the ciHHV-6A in the Japanese individual (HAPMAP NA18999) shows greater divergence from the other ciHHV-6A samples of European origin.

In summary, comparisons of nucleotide substitution frequencies show that the viral genomes in ciHHV-6B carriers are more similar to each other than they are to reference genomes derived from clinical isolates of non-integrated HHV-6B from Japan (HST) and D.R.Congo (Z29). The ciHHV-6A genomes are also more similar to each other than they are to the three HHV-6A reference genomes, although this is less pronounced than seen among the ciHHV-6B genomes.

### Phylogenetic analysis of ciHHV-6 and non-integrated HHV-6 genomes

Consistent with the results shown in Fig. 2, phylogenetic analysis of the U region from 21 ciHHV-6B and the HST and Z29 reference genomes (excluding DR, the large repeat regions and missing data shown in Fig. 1) shows that the ciHHV-6B genomes in HGDP00813 from China and HGDP00092 from Pakistan are outliers to the 19 ciHHV-6B genomes from individuals of European descent (Fig. 3A). A phylogenetic network of the ciHHV-6B genomes with European ancestry shows three clusters of 8, 3 and 5 closely related ciHHV-6B genomes (groups 1, 2 and 3, respectively; Fig. 3B) and three singletons (ORCA1340, COR264 and 1-ciHHV-6B). Phylogenetic analysis of DR alone shows that, with the exception of COR264, the European ciHHV-6B samples show greater similarity to the HST (Japan) reference genome than to the Z29 (D.R.Congo) reference genome. However, the DRs in the two non-European ciHHV-6B samples HGDP000813 (China) and HGDP00092 (Pakistan) do not cluster closely with those in the European ciHHV-6B samples again indicating these ciHHV-6B strains are distinct (Supplementary Fig. S1 and Fig. 3A).

**Figure 3.**
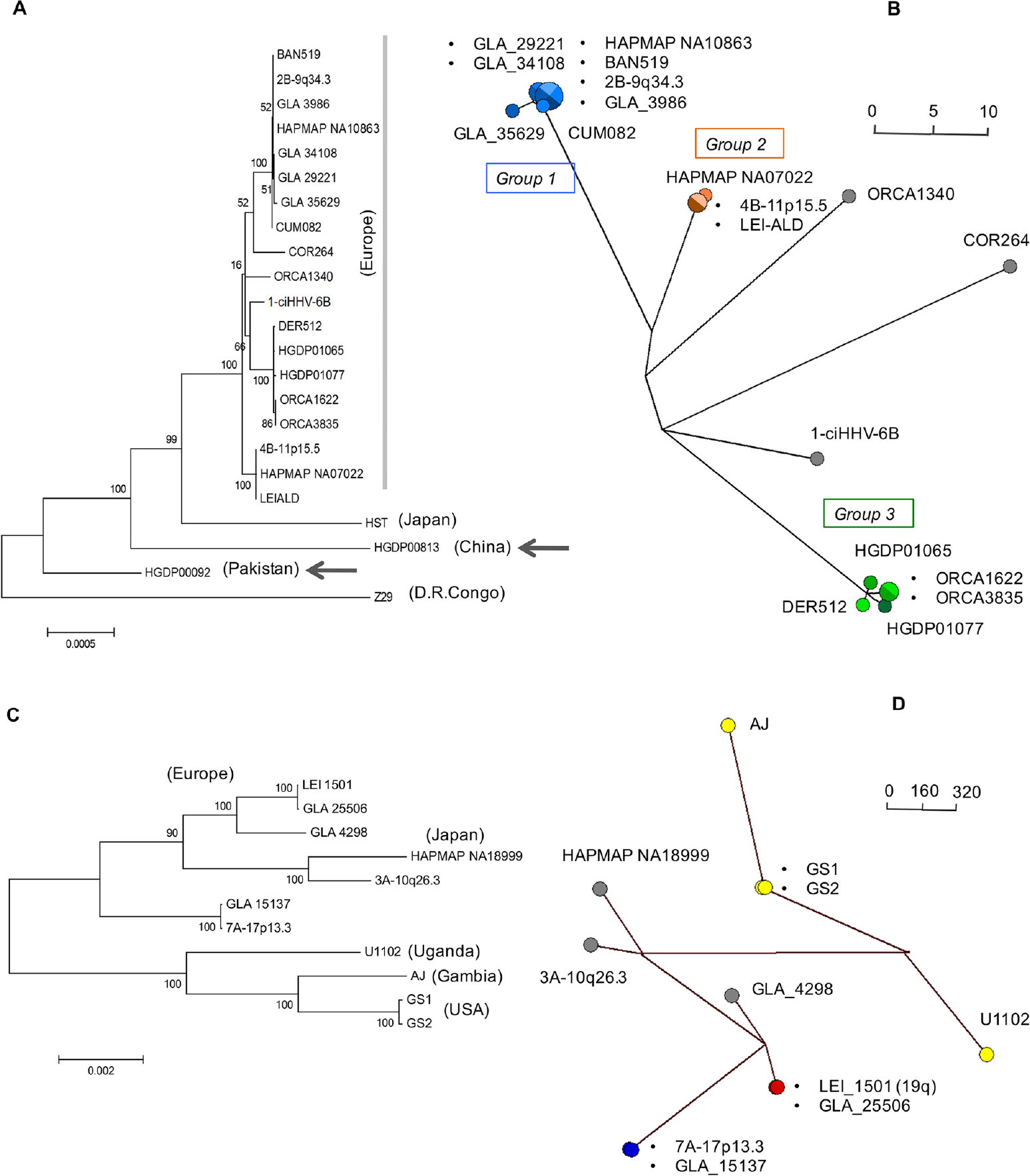
Phylogenetic analysis of ciHHV-6 and reference non-integrated HHV-6 genomes. (A) Maximum likelihood phylogenetic tree of 21 ciHHV-6B genomes and two HHV-6B reference genomes (strains HST (Japan) and Z29 (D.R Congo)). A total of 130412 nucleotides were analysed, excluding repeat regions and missing amplicons. The scale bar represents 0.0005 substitutions per site. (B) Phylogenetic network generated from the dataset used in (A), but without the HST and Z29 genomes and the ciHHV-6B genomes from HGDP00813 (China) and HGDP00092 (Pakistan). The ciHHV-6B genomes from Europeans in groups 1, 2 and 3 are shown as blue, orange and green dots, respectively and the singletons are shown as grey dots. (C) Maximum likelihood phylogenetic tree of seven ciHHV-6A genomes and four HHV-6A reference genomes (strains U1102 (Uganda), AJ (Gambia), GS1 (USA) and GS2 (USA); GS1 and GS2 are two versions of strain GS). A total of 117900 nucleotides were analysed, excluding repeat regions and missing amplicons. The scale bar represents 0.002 substitutions per site. (D) Phylogenetic network generated from the dataset used in (C). The non-integrated HHV-6A reference genomes are shown as yellow dots. The closely related ciHHV-6A genomes are shown as pairs of red or blue dots and singletons as grey dots (including one from Japan). The scale bars in the networks (C and D) show the number of base substitutions for a given line length. The dots are scaled, the smallest dot representing a single individual.

To explore variation only within HHV-6B genes, the frequency of substitutions in ORFs of each of the 21 ciHHV-6B genomes were compared with those in the HST and Z29 reference genomes and the ciHHV-6B genome in HAPMAP NA10863 (Fig. 4A). The patterns of variation were similar to those observed across the whole genome (Fig. 2A) and consistent with the phylogenetic analysis showing greater similarity among ciHHV-6B in Europeans and with the subgroups. Phylogenetic analysis of specific genes, which were selected because they show greater sequence variation from the reference genomes or among the ciHHV-6B genomes, generated a variety of trees that are generally consistent with the phylogenetic analysis based on the U region but exhibited less discrimination between samples or groups (Fig. 4 and Supplementary Fig. S2). For example, the phylogenetic tree based on U90 separates the European ciHHV-6B samples from the ciHHV-6B samples from China and Pakistan and from the HST and Z29 reference genomes but does not subdivide the European ciHHV-6B samples.

**Figure 4.**
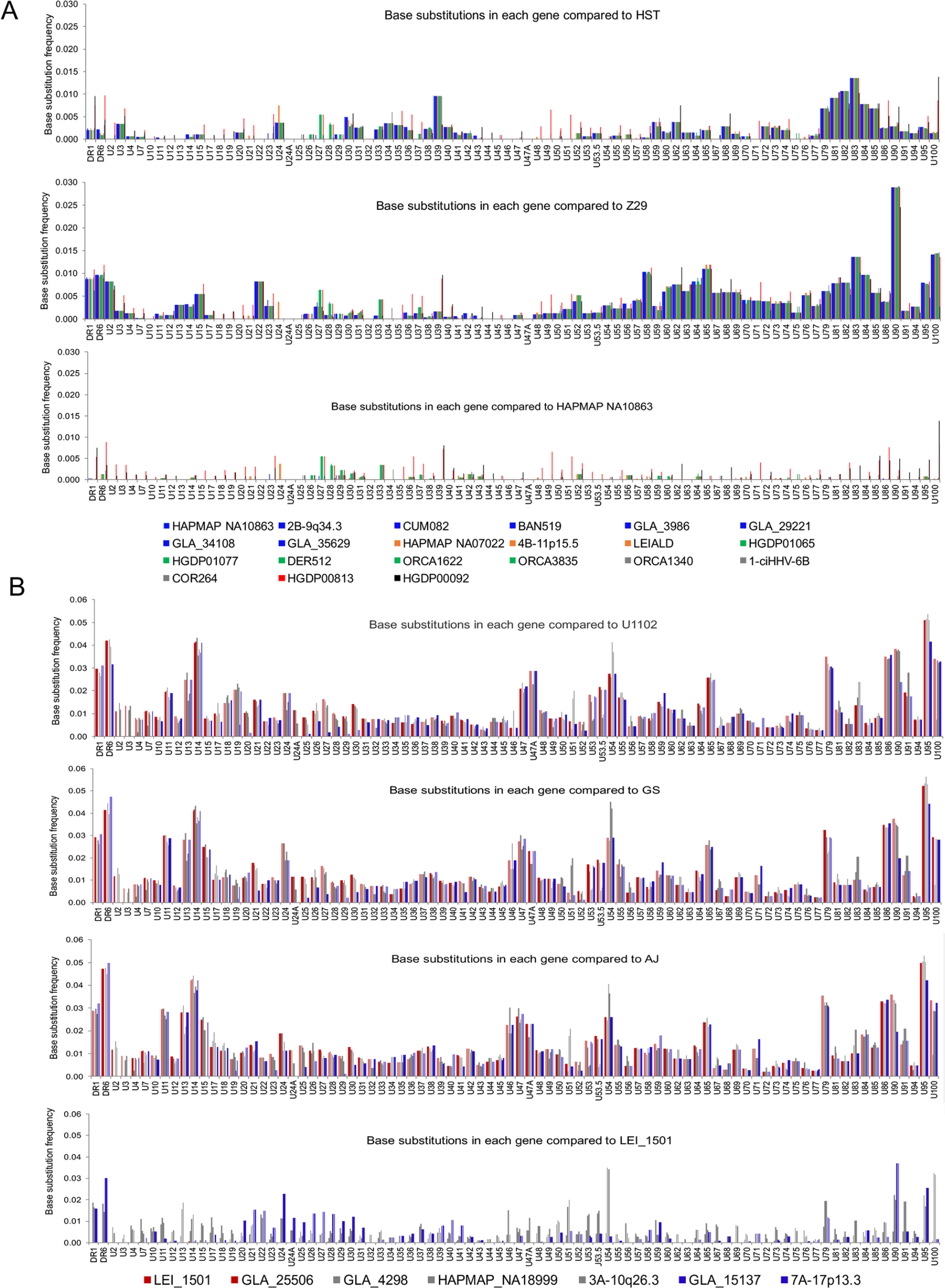
Frequency of nucleotide substitutions in ciHHV-6 genes compared to those in reference viral genomes. (A) Graphs of substitution frequency in each gene is shown for the 21 ciHHV-6B genomes in comparison with HHV-6B strains HST (Japan) and Z29 (D.R.Congo) genomes (top and middle panels, respectively) and the ciHHV-6B genome in European HAPMAP NA10863 (bottom panel). The colour coding shown in the key matches that of the network in Figure 3B as follows: European Group 1, pale blue lines; European Group 2, orange lines; European Group 3, green lines; European singletons, grey lines; ciHHV-6B in HGDP00813 from China, red lines; and ciHHV-6B in HGDP00092 from Pakistan, black lines. (B) Graphs of substitution frequency in each gene for each of the 7 ciHHV-6A genomes in comparison with the HHV-6A strains U1102 (Uganda), GS (USA) and AJ (Gambia) genomes (top and two middle panels) and the ciHHV-6A genome in European LEI-1501 (bottom panel). The colour-coded key matches that of the network in Figure 3D. The x-axes of all the graphs show a single copy of DR1 and DR6, followed by genes found in the U region.

Phylogenetic analysis of the seven ciHHV-6A genomes and four reference genomes (U1102 (Uganda), AJ (Gambia) and two sequences from GS (USA)) shows a clear separation between the integrated and non-integrated genomes (Fig. 3C and D), with two pairs of closely related ciHHV-6A genomes (LEI-1501 and GLA_25506; 7A-17p13.3 and GLA_15137). A similar separation of the integrated versus non-integrated genomes is also evident in the phylogenetic analysis of DR alone, irrespective of the geographic origin of the individual ciHHV-6A carrier (Supplementary Fig. S1).

Variation within HHV-6A genes was also explored by plotting base substitution frequency per ORF for each of the seven ciHHV-6A samples in comparison to the three reference genomes and the ciHHV-6A genome in LEI_1501 (Fig. 4B). The patterns of variation are similar to those observed across the whole genome (Fig. 2B). Phylogenetic analysis of U83, U90 and DR6, selected because they show greater sequence variation, generally support the phylogenetic trees and networks generated from analysis of the Uand DR regions (Supplementary Fig. S3).

Overall, the sequence variation and phylogenetic analyses indicate a divergence between the integrated and non-integrated HHV-6 genomes but with some differences between the HHV-6A and HHV-6B. The ciHHV-6B samples from individuals with European ancestry showed divergence from both HST (Japan) and Z29 (D.R.Congo) reference genomes although the pattern of divergence varies across the genome. The 21 ciHHV-6B genomes from individuals with European ancestry are more similar to one another than to the ciHHV-6B genomes from China and Pakistan and can be subdivided into distinct groups. There is greater divergence among the seven ciHHV-6A genomes than among the ciHHV-6B genomes but despite this two pairs of closely related ciHHV-6A genomes were identified.

From these analyses we concluded that the three groups of closely related ciHHV-6B genomes and the pairs of ciHHV-6A genomes identified in the phylogenetic networks (Fig. 3B and D, respectively) could represent independent integrations by closely related strains of HHV-6B or HHV-6A. Alternatively, each group might have arisen from a single integration event, with members sharing a common ancestor. Further analyses were undertaken to explore these possibilities.

### Comparison of tandem repeat regions in ciHHV-6 genomes

Tandem repeat arrays within the human genome often show length variation as a consequence of changes to the number of repeat units present (copy number variation). The greater allelic diversity in these regions reflects the underlying replication-dependent mutation processes in tandem repeat arrays, which occur at a higher rate than base substitutions (39). To explore diversity among the ciHHV-6B genomes further, tandem repeat regions distributed across the viral genome were investigated. The R-DR, R2A, R2B and R4 repeat regions analysed (location shown in Fig. 1C) showed little or no copy number variation among the ciHHV-6B and non-integrated reference genomes (Fig. 5A, Table 2). Copy number variation at R1 (location shown in Fig. 1C) was greater but did not show a clear relationship with strains of ciHHV-6B or non-integrated HHV-6B. Greater copy number variation was detected at the pure array of TTAGGG repeats at DR_L_-T2 (location shown in Fig. 5B) with the largest number of repeats in the HHV-6B Z29 reference genome and ciHHV-6B in HGDP00813 from China (Fig. 5A, Table 2). Notably, copy number variation observed at R0 (location shown in Fig. 1C) correlates reasonably well with the groups of ciHHV-6B genomes identified the phylogenetic network (Fig 5A; Table 2; Fig. 3).

**Figure 5.**
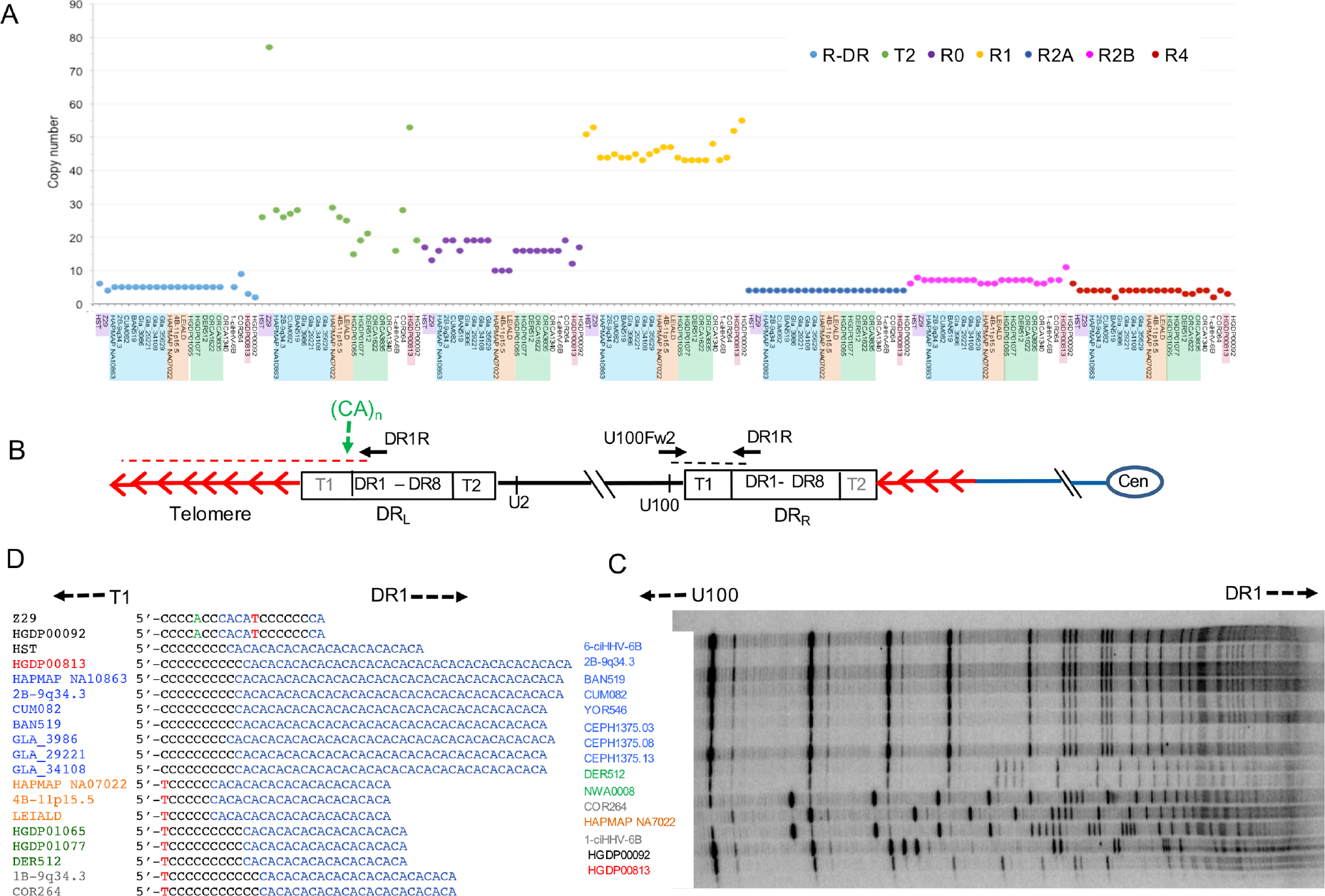
Copy number variation in tandem repeat loci across the HHV-6B genome. (A) Graph of the number of repeat units at loci within the DR (R-DR and DR_L_-T2) and U regions (R0, R1, R2A, R2B and R4). Comparisons can be made among the reference non-integrated HHV-6B strains, HST (Japan) and Z29 (D.R. Congo) and ciHHV-6B genomes. The sample order along the x-axis as follows: HST, Z29 (mauve highlight); European group 1 ciHHV-6B genomes (blue highlight); European group 2 ciHHV-6B genomes (orange highlight); European group 3 ciHHV-6B genomes (green highlight); European singleton ciHHV-6B genomes (no highlight); ciHHV-6B in HGDP00813 from China (red highlight); and ciHHV-6B in HGDP00092 from Pakistan (no highlight). (B) Diagram showing the location of the PCR amplicons used to analyse the repeat sequences shown in C and D. Black dashed line shows the amplicon generated by the U100Fw2 and DR1R primers that were used for TVR-PCR shown in (C). Red dashed line shows STELA products, generated from DR1R, that were used to analysis the (CA)_n_ repeat shown in D. (C) Distribution of (TTAGGG) repeats at the distal end of DR_R_-T1 (near U100) in ciHHV-6B genomes. Each black rung in the ladder shows the position of a (TTAGGG) repeat, the intervening sequence longer that 6bp comprise degenerate repeats. The ciHHV-6B sample names are colour-coded in accordance with groupings identified in Figure 3: European group 1, blue; group 2, orange; group 3, green; European singletons, grey; ciHHV-6B in HGDP00813 from China, red; ciHHV-6B in HGDP00092 from Pakistan, black. (D) Variation in copy number of (CA) repeats and adjacent 5’-sequence, near the start of the ciHHV-6B DR_L_-T1 region. Sample names colour-coded as described in (C).

**Table 2.** Variation in tandem repeat regions among ciHHV-6.

| Tandem repeat regions in HHV-6B |  |  |  |  |  |  |  |  |  |  |
| --- | --- | --- | --- | --- | --- | --- | --- | --- | --- | --- |
| Name | T1 | STR (CA)<sub>n</sub> | R-DR<sub>a</sub> | T2 | R0<sup>a</sup> | R1 | R2A<sub>a</sub> | R2B | R3 | R4<sup>a</sup> |
| Location |  | adjacent to DR-T1 | DR | DR | U1 | U86 | U86-U89 | U86-U89 | U91-U94 | After U100 |
| Length (bp) | 6 | 2 | 15 | 6 | ~15 | 12 | 79 | 12 - 15 | ~104 | 64 |
| Unit |  | CA | NI<sup>b</sup> | (TTAGGG) | NI | NI | NI | NI | NI | NI |
| **HST<sup>c</sup>** | - | 12 | 6 | 26 | 17 | 51 | 4 | 6 | - | 6 |
| **Z29<sup>c</sup>** | - | 1 | 4 | 77 | 13 | 53 | 4 | 8 | - | 4 |
| HAPMAP NA10863 | - | 20 | 5 | 28 | 16 | 44 | 4 | 7 | - | 4 |
| 2B-9q34.3 | - | 20 | 5 | 26 | 19 | 44 | 4 | 7 | - | 4 |
| CUM082 | - | 19 | 5 | 27 | 19 | 45 | 4 | 7 | - | 4 |
| BAN519 | - | 19 | 5 | 28 | 16 | 44 | 4 | 7 | - | 4 |
| GLA_3986 | - | 20 | 5 | - | 19 | 44 | 4 | 7 | - | 2 |
| GLA_29221 | - | 19 | 5 | - | 19 | 45 | 4 | 7 | - | 4 |
| GLA_34108 | - | 19 | 5 | - | 19 | 43 | 4 | 7 | - | 4 |
| GLA_35629 | - | - | 5 | - | 19 | 45 | 4 | 7 | - | 4 |
| HAPMAP NA07022 | - | 11 | 5 | 29 | 10 | 46 | 4 | 6 | - | 4 |
| 4B-11p15.5 | - | 11 | 5 | 26 | 10 | 47 | 4 | 6 | - | 4 |
| LEI-ALD | - | 11 | 5 | 25 | 10 | 47 | 4 | 6 | - | 4 |
| HGDP01065 | - | 10 | 5 | 15 | 16 | 44 | 4 | 7 | - | 4 |
| HGDP01077 | - | 10 | 5 | 19 | 16 | 43 | 4 | 7 | - | 4 |
| DER512 | - | 10 | 5 | 21 | 16 | 43 | 4 | 7 | - | 4 |
| ORCA1622 | - | - | 5 | - | 16 | 43 | 4 | 7 | - | 3 |
| ORCA3835 | - | - | 5 | - | 16 | 43 | 4 | 7 | - | 3 |
| ORCA1340 | - | - | - | - | 16 | 48 | 4 | 6 | - | 4 |
| 1-ciHHV-6B | - | 12 | 5 | 16 | 16 | 43 | 4 | 6 | - | 4 |
| COR264 | - | 12 | 9 | 28 | 19 | 44 | 4 | 7 | - | 2 |
| HGDP00813 | - | 20 | 3 | 53 | 12 | 52 | 4 | 7 | - | 4 |
| HGDP00092 | - | 1 | 2 | 19 | 17 | 55 | 4 | 11 | - | 3 |
| Tandem repeat regions in HHV-6A |  |  |  |  |  |  |  |  |  |  |
|  | T1 | T2 | R5<sup>d</sup> | R1 | R2 | R3 |  |  |  |  |
| Location |  | DR | U41 - U42 | U86 | U86 - U89 | U91 - U94 |  |  |  |  |
| Length (bp) | 6 | 6 | ~191 | ~12 | 12-18 | 104-105 |  |  |  |  |
|  |  | (TTAGGG) | NI | NI | NI | NI |  |  |  |  |
| **AJ<sup>c</sup>** | - | 51 | 1.7 | 52 | 43 | 8 |  |  |  |  |
| **U1102<sup>c</sup>** | - | 59 | 1.7 | 52 | 102 | 29 |  |  |  |  |
| **GS1/2<sup>c</sup>** | - | 51 | 1.7 | 52 | 78 | 8 |  |  |  |  |
| LEI-1501 | - | 14 | 2.7 | - | - | - |  |  |  |  |
| GLA_25506 | - | - | 2.7 | 32 | - | - |  |  |  |  |
| GLA_4298 | - | - | 3.7 | 53 | - | - |  |  |  |  |
| HAPMAP NA18999 | - | 13 | 1.7 | - | - | - |  |  |  |  |
| 3A-10q26.3 | - | 9 | 1.7 | 58 | - | - |  |  |  |  |
| GLA_15137 | - | - | 1.7 | 55 | - | - |  |  |  |  |
| 7A-17p13.3 | - | - | 1.7 | 55 | - | - |  |  |  |  |

<sup>a</sup> Repeats specific to HHV-6B - the coordinates of R-DR and R4 in HHV-6B strain HST are 5400-5489 and 152603-152986 respectively; <sup>b</sup> NI, repeats not identical; <sup>c</sup> Reference genomes in bold; <sup>d</sup> Repeat specific to HHV-6A - the coordinates of R5 in HHV-6A strain U1102 are 68124-68450; the other repeats are described in (9) (25); hyphens, analysis not completed. The samples in the same box are in the same group in the phylogenetic networks.

Similar analysis of repeat regions in the ciHHV-6A genomes was conducted (Table 2). The data suggest that ciHHV-6A genomes have fewer TTAGGG repeats at DR_L_-T2 than in the HHV-6A reference genomes. This variation could have been present in HHV-6A strains prior to integration or deletion mutations that reduce the length of the DR_L_-T2 array may have been favoured after integration (12).

To explore variation within the T1 array of degenerate telomere-like repeats in ciHHV-6B genomes we amplified DR_R_-T1 region using the U100Fw2 and DR1R primers and investigated the interspersion patterns of TTAGGG and degenerate repeats at the distal end of DR_R_-T1 (near U100, Fig. 5B) using modified TVR-PCR (35, 36, 40). The ciHHV-6B genomes clustered into groups that share similar TVR maps in DR_R_-T1 (Fig. 5C) and the patterns differed among the groups and the singleton ciHHV-6B genomes identified in the phylogenetic analyses. Variation around the (CA)_n_ simple tandem repeat, located immediately adjacent to DR_L_-T1 (location shown in Fig. 5B), also showed clustering into groups that correlate with the ciHHV-6B phylogenetic analyses (Fig 5D, Table 2, Fig. 3). Overall, the analyses of tandem repeat regions in the ciHHV-6B genomes are consistent with the phylogenetic analyses.

### Ancestry of ciHHV-6B carriers in group 3

The repeat copy number variation observed within and among groups may have arisen before or after telomeric integration of the viral genome. To investigate further how many different integration events may have occurred among the ciHHV-6B carriers, we isolated and sequenced fragments containing the junction between the human chromosome and the ciHHV-6B genome, in addition to using the cytogenetic locations published previously for some samples (Table 1; (16)). The junction fragments were isolated by a trial-and-error approach, using PCR between a primer mapping in DR8 in DR_R_ and a variety of primers known to anneal to different subtelomeric sequences (Fig. 6A), including primers that anneal to the subterminal region of some but not all copies of chromosome 17p (17p311 (41) and subT17-539 (12)). There was insufficient DNA for analysis from the sequenced ORCA1340 (singleton) or the ORCA1622 and ORCA3835 (group 3) samples (Fig. 3B). However, analysis of DR_R_-T1 and the other repeats showed that the 42 ciHHV-6B carriers from Orkney fall into two groups, that share the same length at DR_R_-T1 with either ORCA1340 or with ORCA1622 and ORCA3835 (Table 2). For junction fragment analysis, we selected ORCA1006 as a substitute for ORCA1340, since it shares the same DR_R_-T1 length. Similarly, ORCA1043, ORCA2119 and ORCA1263 were used as substitutes for ORCA1622 and ORCA3835, since they share a different DR_R_-TI length. Using the chromosome 17p primers, junction fragments were generated from all the group 3 ciHHV-6B samples and from 1-ciHHV-6B (a singleton in the phylogenetic network, Fig. 3). PCR products were not amplified using these primers from other ciHHV-6B samples in this study or from another sample with a reported location at 17p (5B-17p13.3 (16), data not shown). The sequences of seven junction fragments from group 3 ciHHV-6B genomes (including NWA008 (42), which is another ciHHV-6B carrier having a viral genome that belongs to group 3 (data not shown)) were similar to each other but different from the fragment in sample 1-ciHHV-6B (Fig. 6B). These data indicate the existence of three independent integration events into different alleles of the chromosome 17p telomere, or possibly into telomeres of different chromosomes that share similar subterminal sequences (43).

**Figure 6.**
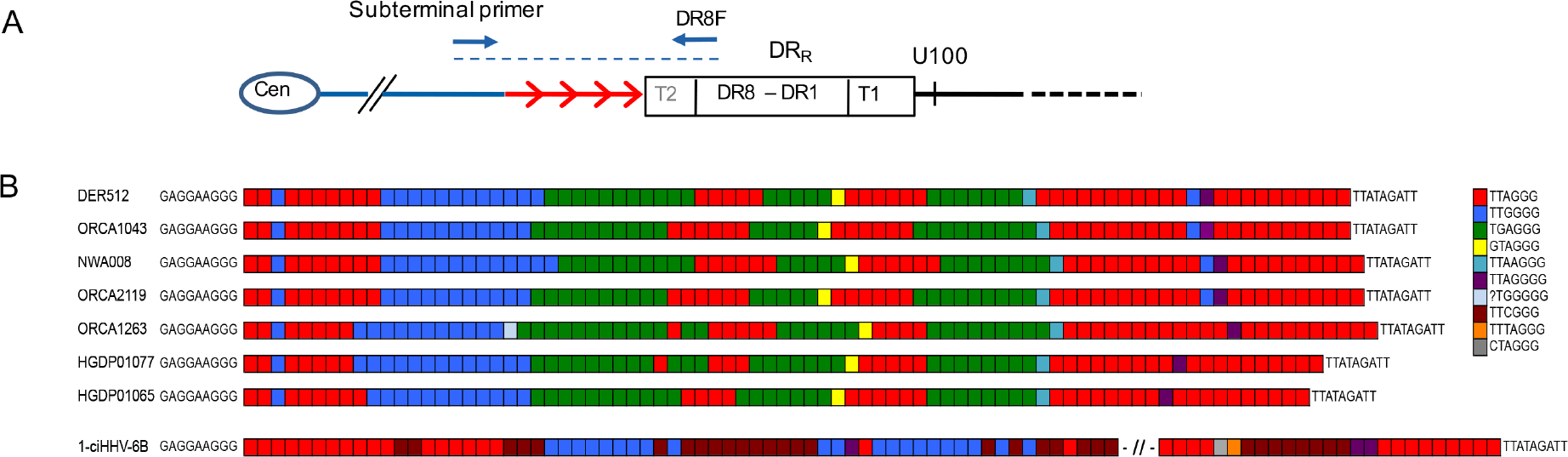
Characterisation of ciHHV-6B integration sites. (A) Diagram showing the location of PCR amplicons used to characterise the chromosome-ciHHV-6B junctions. Red arrows represent TTAGGG and degenerate repeats. Blue arrows, primers used to amplify the chromosome-HHV-6 junction; blue dashed line, chromosome-junction amplicon used for sequence analysis. (B) Diagram showing the similarity of the TTAGGG (red squares) and degenerate repeat (coloured squares in key to right) interspersion patterns in the chromosome-HHV-6 junctions from individuals with group 3 ciHHV-6B genomes (DER512 to HGDP01065, Figure 3B). These interspersion patterns are distinct from that of the chromosome-junction fragment isolate from 1-ciHHV-6B (singleton in Figure 3B). The sequence to the left of the repeats is from the chromosome subtelomeric region and the sequence to the right is from the ciHHV-6B genome.

Comparison of the junction fragments from group 3 ciHHV-6B samples shows remarkably similar TTAGGG and degenerate repeat interspersion patterns (Fig. 6B). The differences among the interspersion patterns are consistent with small gains or losses that may have arisen from replication errors in the germline, after integration of the viral genome (35). Therefore, it is most likely that the ciHHV-6B status of group 3 individuals arose from a single ancestral integration event. Using the levels of nucleotide substitution between the group 3 ciHHV-6B genomes, the time to the most recent common ancestor (TMRCA) was estimated as 24,538 ±10,625 years ago (Table 3). This estimate is based on the assumption that once integrated the ciHHV-6B genome mutates at the same average rate for the human genome however, deviation from this would result in an under or over estimation of the TMRCA.

**Table 3.** Estimate of TMRCA for ciHHV-6B genomes in group 3.

|  | Entire group 3 <sup>a</sup> | HGDP1065 &<br>HGDP1077 | HGDP1065 &<br>DER512 |
| --- | --- | --- | --- |
| TMRCA (y) | 24,538 | 23,004 | 15,336 |
| Standard deviation | 10,625 | 13,281 | 10,844 |

<sup>a</sup> ORCA1622 and ORCA3835 are identical across non-repeat regions.

### Genetic intactness of ciHHV-6 genomes

The evidence for an ancient origin of some, probably most, of the ciHHV-6B genomes analysed, and for post-integration mutations in repeat regions raised the question of whether these genomes contain an intact set of viral genes or whether they have been rendered non-functional by mutation. To explore the consequence of sequence variation among the ciHHV6B genomes, the amino acid sequences predicted from all genes in the ciHHV-6B genomes were aligned, and the cumulative frequencies of independent synonymous and non-synonymous substitutions were determined (Fig. 7A). The ratio of synonymous:non-synonymous substitutions varies among genes. The great majority of non-synonymous changes (amounting to 34% of the total) result in single amino acid substitutions, but one substitution in the U20 stop codon of HGDP00092 is predicted to extend the coding region by eight codons. Only one substitution, which creates an in-frame stop codon in U14 of 1-ciHHV-6B, is predicted to terminate a coding region prematurely. Two of the seven ciHHV-6A genomes also have in-frame stop codons, one in U79 of GLA_15137 and the other in U83 genes of GLA_4298 (data not shown).

**Figure 7.**
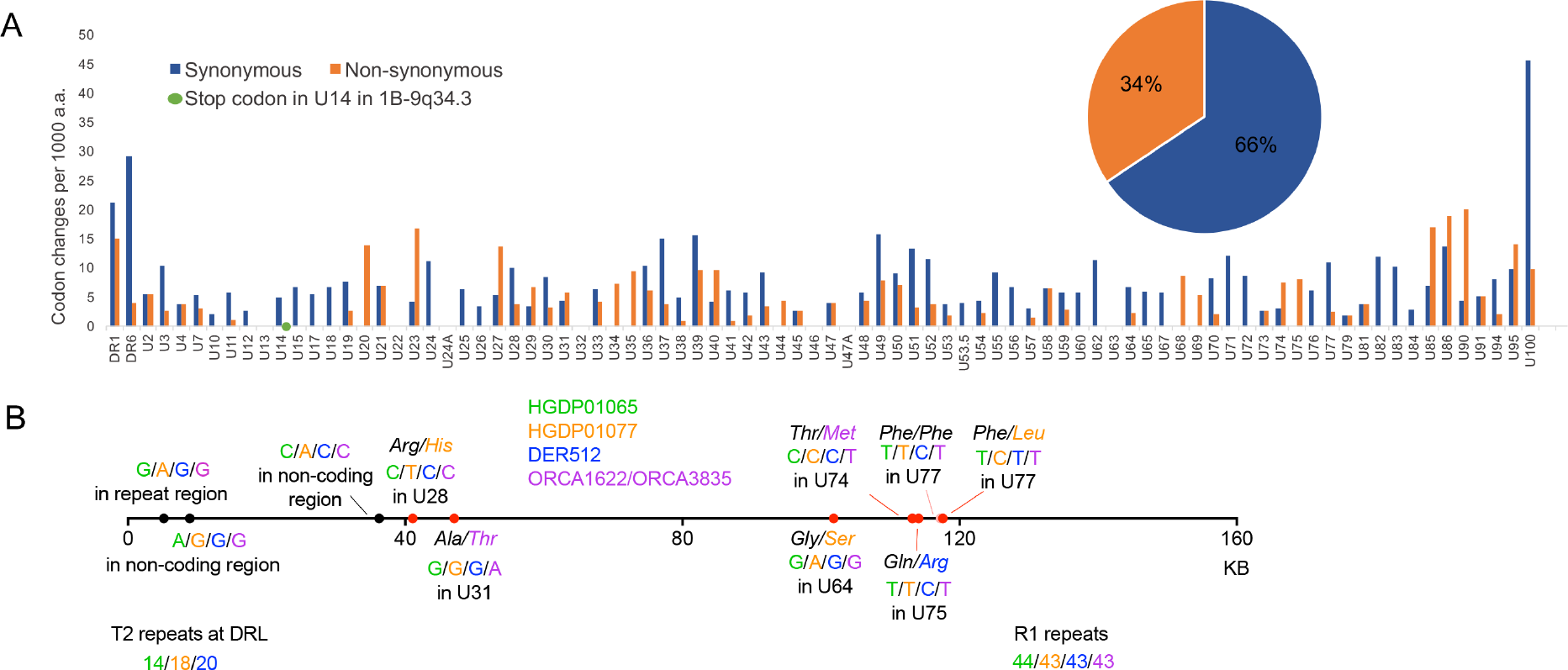
Consequences of nucleotide substitutions across the ciHHV-6 genome. (A) Comparison of synonymous (blue) and non-synonymous (orange) substitution frequencies in each ciHHV-6B gene among the 21 ciHHV-6B genomes (scaled to differences per 1000 amino acids). The green dot shows the novel in-frame stop codon in U14 of 1-ciHHV-6B. The pie chart shows the overall proportions of synonymous and non-synonymous substitutions across all genes. (B) Diagram showing the approximate location and consequence of nucleotide substitutions that are predicted to have arisen after integration in group 3 ciHHV-6B genomes. The horizontal line represents the HHV-6B genome; black dots, location of non-coding base substitutions; red dots, base substitutions within HHV-6B genes that are predicted to result in an amino acid substitutions (non-synonymous) shown by the text; pink dot, synonymous (T to C) substitution in DER512 that is not predicted to change the phenylalanine. HGDP01065, green text; HGDP01077, orange text; DER512 in blue and the identical sequences found in ORCA1622 and ORCA3835 in purple. The number of repeats in three regions (T2, R1 and R4) that vary among the group 3 genomes are also shown.

The 21 iciHHV-6B genomes are likely to include mutations that arose before integration and represent variation among the parental non-integrated HHV-6B strains, as well as mutations that arose after integration. To explore the latter, five group 3 ciHHV-6B genomes were compared (Fig. 7B). Among the ten substitutions identified, three were in non-coding regions, one was a synonymous mutation in U77, and six were non-synonymous mutations. From these limited data, it seems likely that the accumulation of mutations after integration has been random in these ciHHV-6B genomes.

## DISCUSSION

In this study we have used comparative analyses to explore diversity among ciHHV-6 genomes, to understand the factors that influence the population frequencies of ciHHV-6 and to determine whether the integrated genomes retain the capacity for full functionality as a virus. We have found that the ciHHV-6B genomes are more similar to each another than to the two available HHV-6B reference genomes from Japan and D.R.Congo (Figs 2, 3, 4; Supplementary Fig. S1). This is particularly noticeable among the 19 ciHHV-6B genomes from individuals with European ancestry, which are more similar to each other than they are to the ciHHV-6B genomes in HGDP00092 from Pakistan and HGDP00813 from China. This pointer towards a relationship between the integrated HHV-6B strain and geographical distribution warrants further investigation, if the association between carrier status and potential disease risk is to be understood fully (21, 44). The smaller group of seven ciHHV-6A genomes show higher levels of divergence from the three available HHV-6A reference genomes from the USA, Uganda and the Gambia. However, in making these observations, the possibility of sample bias should be considered, both in the geographic distribution of ciHHV-6 genomes analysed and, in particular, in the small number of non-integrated HHV-6A and HHV-6B genomes that are available for comparative analysis.

The isolation of chromosome junction fragments from eight ciHHV-6B samples (seven group 3 samples and 1-ciHHV-6B) by using primers from chromosome 17p subterminal sequences (41) suggests integration in alleles of the 17p telomere. Given the variable nature of human subterminal regions (43), the chromosome locations should be confirmed using a different approach. Nevertheless, comparison of the TTAGGG and degenerate repeat interspersion patterns at the chromosome-ciHHV-6B junction can be used to deduce relationships (40, 45) and, combined with the phylogenetic analyses, show that the individuals carrying a group 3 ciHHV-6B genome share an ancient ancestor. Group 3 includes individuals from Sardinia, England, Wales and Orkney, with greater divergence between the ciHHV-6B genomes in the two individuals from Sardinia (HGDP1065 and HGDP1077) than between the individual from Derby, England (DER512) and the Sardinian (HGDP1065) (Figs 3, 5, 6 and Tables 2, 3). Moreover there is no evidence of a close family relationship between the two individuals from Sardinia. Overall the data are consistent with the group 3 ciHHV-6B carriers being descendants of a common ancestor who existed approximately 24,500 years ago, similar to the date of the last glacial maximum and probably predating the colonization of Sardinia and Orkney.

The population screen of Orkney identified 42 ciHHV-6B carriers (frequency 1.9%, Supplementary Table S1) and no ciHHV-6A carriers, which also suggests a founder effect. However, the Orkney ciHHV-6B samples can be divided into two groups, based on the length of DR_R_-T1, the ciHHV-6B phylogenetic analyses and the different integration sites. Therefore, it is likely that the ciHHV-6B carriers in Orkney are the descendants of two different ciHHV-6B ancestors, who may have migrated to Orkney independently. This is consistent with the fine resolution genetic structure of the Orkney population and the history of Orkney, which includes recent admixture from Norway (Norse-Vikings) (42).

Given the evidence that extant ciHHV-6B carriers in group 3 are descendants of a single ancient founder with a germline integration, it is plausible that other clusters in the phylogenetic tree have a similar history. For example the three individuals in group 2 may all carry a ciHHV-6B integrated in a chromosome 11p telomere. Further verification is required to support this speculation, and this will be valuable when assessing disease risk associated with ciHHV-6 integrations in different telomeres.

There is good evidence that ciHHV-6 genomes can reactivate in some settings, for example when the immune system is compromised (19, 20). However, it is not known what proportion of ciHHV-6 genomes retain the capacity to reactivate. We investigated this question from various angles. We presented evidence that some ciHHV-6 genomes are ancient and therefore could have accumulated inactivating mutations while in the human genome. Most of the tandem repeats analysed in ciHHV-6B genomes showed minor variations in repeat copy numbers (Fig. 5 and Table 2). However, the function of these regions is unclear, and as copy number variation exits among the reference genomes, it is unlikely that the level of variation detected unduly influences the functionality of the viral genome. In the protein-coding regions of ciHHV-6B genomes, 34% of substitutions are non-synonymous and are predicted to cause amino acid substitutions (Fig. 7). A single potentially inactivating mutation was detected as an in-frame stop codon in gene U14 in 1-ciHHV-6B. Since this gene encodes a tegument protein that is essential for the production of viral particles and can induce cell cycle arrest at the G2/M phase (46), it seems unlikely that this integrated copy of ciHHV-6B would be able to reactivate. However, the other viral genes may be expressed in this ciHHV-6B genome and the presence of the viral genome may also affect telomere function. The stop codon in gene U20 in the individual from Pakistan (HGDP00092) is mutated, and this is predicted to extend the U20 protein by eight amino acid residues. U20 is part of a cluster of genes (U20-U24) that are specific to HHV-6A, HHV-6B and their relative human herpesvirus 7, and likely plays a role in suppressing an apoptotic response by the infected host cell (47, 48). Further experimental analysis will be required to determine whether the extension affects the function of the U20 protein. Among the seven ciHHV-6A genomes, two contain novel in-frame stop codons. One of these is located in U83 in GLA_4298. The other is present in U79 in GLA_15137, but this inactivating mutation is absent from the closely related ciHHV-6A genome in 7A-17p13.3 (Fig. 3C and D).

In summary, we have shown that most ciHHV-6A and ciHHV-6B genomes contain an intact set of genes and therefore may have the potential to be fully functional. This observation needs to be taken into consideration when assessing whether ciHHV-6 carrier status is associated with disease risk and in understanding the underlying mechanisms of such associations (e.g. whether viral reactivation is involved). Among the individuals of European descent, we found strong evidence for the ancient common ancestry of some of the integrated viral genomes. The close similarity between ciHHV-6B genomes in the Europeans and the evidence of multiple different integration events by similar strains also indicate that we have effectively sequenced the ancient, non-integrated strains of HHV-6B that existed in European populations in prehistoric times. Based on these observations, it is possible that other populations, for example in China, South Asia and Africa, may show similar founder effects among ciHHV-6 carriers but from different ancient strains. Our limited knowledge of non-integrated HHV-6A and HHV-6B strains is based mostly on strains derived from Africa and Japan. There is now a real need to sequence non-integrated strains from other populations, including those in Europe, so that the relationship between non-integrated HHV-6 and ciHHV-6 can be fully understood. A major challenge will be to determine whether germline integration continues to occur *de novo* today, and, if so, at what rate and by which viral strains.

## ACCESSION NUMBERS

The finished sequences have been deposited in GenBank under accession numbers KY316030-KY316056. The LEI_1501 ciHHV-6A genome reported previously has the accession number KT355575 (18).

## SUPPLEMENTARY MATERIAL

Supplementary data are available online.

## ACKNOWLEDGMENTS

We thank most sincerely Mark Jobling, Michael Wood and Ryan Mate (University of Leicester) for their help with data analysis. We also thank Martin Dyer (University of Leicester), Bruce Winney (University of Oxford), James F. Wilson (University of Edinburgh) and Duncan A. Clark (Department of Virology, Barts Health NHS Trust) for samples from the various populations screened. Author contributions: EZ, AJB, GW, NMS, RN, IIAC, VEC and YH conducted various aspects of the experimental work; AD, EZ, GW, NMS, CV and CB conducted the bioinformatic and other analyses; DJP is a member of the Executive Committee of Generation Scotland and, with AJB and RFJ, was involved in the identification of ciHHV-6 samples from the GS: SFHS cohort; NJR was responsible for the project design. The paper was written by EZ and NJR with significant input from AD and RFJ. All authors reviewed the manuscript.

